# Genome-Wide Identification of G-Quadruplex forming regions in Arabidopsis: Unraveling the Role of Pif like Helicase (AtPLH1) in Gene Regulation and Stress Response

**DOI:** 10.1101/2024.03.11.584348

**Authors:** Surabhi Singh, Shubhangi Singh, Khushboo Gupta, Himanshi Sharma, Shivsam Bady, Manushka Sondhi, Rohini Garg

**Author notes:** Corresponding author: Rohini Garg. Equal contribution (Surabhi Singh, Shubhangi Singh, Khushboo Gupta).

## Abstract

G-quadruplexes (GQSes) are highly stable DNA secondary structures, which exist as knots in the genome during different cellular processes like replication, transcription and translation. Although several studies have shown the role of GQS-helicases regulating several cellular processes in yeast and human, their detailed characterization in plants is still lacking. In this study, we identified GQS-enriched regions by DNA affinity purification followed by sequencing from Arabidopsis Pif-like helicase 1 (*Atplh1*) mutant. Differentially enriched peaks (DEPs) in the mutant showed preferential distribution in the exonic and promoter regions. The genes involved in various processes like transcriptional regulation, UDP- glucosylation, response to abiotic stress, ethylene biosynthesis and response to carbohydrate stimulus, were found to be differentially regulated between control and mutant plants. The differentially expressed genes (DEGs) showed enrichment of binding sites of ERF, WRKY, BBM and BIM transcription factors. Further, the DEGs harboring DEPs were found associated with response to wounding and salt stress, response to unfolded protein, heat stress response and UDP-glycosyl transferase activity. In addition, the mutants exhibited lesser growth inhibition under cold stress. Overall, our study identified genome-wide GQSes in *Arabidopsis* and altered gene expression regulated by AtPLH1.

## Introduction

In addition to the canonical B-form, DNA can adopt several alternative structures, as commanded by its sequence. These non-canonical structures are referred to as secondary structures, non-B-DNA or alternative DNA and embrace slipped structures, such as Z-motifs, triplex, inverted repeats, mirror repeats, and direct repeats (Sharma et.al., 2011). G- quadruplexes (GQSes) are non-canonical DNA secondary structures arising from the stable folding of single-stranded guanine-rich-DNA sequences. Four guanine bases arranged in a planar quartet interact with each other via Hoogsteen hydrogen bonds form G-quartets, which upon stacking give rise to a GQS and are further stabilized by the presence of monovalent cations, such as K^+^, Na^+^ or Li^+^. Depending on the number of DNA strands contributing towards the formation of GQSes, these can either be unimolecular structures (arising from a single DNA strand) or multimolecular structures (arising from multiple DNA strands).

GQSes play an important role in various cellular processes, such as replication, transcription, and translation. G4 motifs are enriched in the telomeres, promoters and intronic region of genes in various organisms (Huppert & Balasubramanian et al., 2005; Marsico et al., 2019). Although several studies have predicted the genome-wide presence of GQSes across organisms, *in vivo* and *in vitro* validation of GQS formation in these predicted sequences is required (Varshney et al., 2020). Several methods, including Circular Dichroism (CD) spectroscopy, polyacrylamide gel electrophoresis, enzyme mobility shift (assay to identify GQS interacting proteins), DMS footprinting and SHAPE that involves selective 2′- OH acylation analyzed by primer extension etc. are used to identify and characterize GQSes (Han et al., 1999; Dexheimer et al., 2006; Wilkinson et al., 2006; Carey et al., 2012; Lago et al., 2017). Several experimental studies have shown that the G4 motifs are highly enriched in the promoter regions in the genome, suggesting that they might play an important role in gene regulation (Huppert & Balasubramanian et al., 2005). Antibodies highly specific for G-quadruplex structure, such as BG4 and 1H6, have been used for the detection and/or visualization of the DNA GQSes in human and murine cells (Biffi et al., 2013; Henderson *et al*., 2013; Hänsel-Hertsch et al., 2018).

DNA polymerase stop assay, along with G4 sequencing in the presence of G4 stabilizing ligands, have helped in the genome-wide mapping of GQSes (Han et al.,1999; Chambers et al., 2015; Garg et al., 2016). ChIP-qPCR studies in *Saccharomyces cerevisiae* proved pausing of the replication machinery on encountering GQSes in Pif1 helicase mutants (Paeschke et al., 2011). In cellular imaging of GQSes using GQS specific ligands like N-TASQ, DOTASQ etc. or GQS specific antibodies have been used for direct visualization of GQSes (Stefan et al., 2011; Biffi et al., 2013; Henderson et al., 2013; Laguerre et al., 2016). Studies on GQSes interacting proteins like sheltrin complex proteins, nucleolins and GQSes unwinding helicase proteins have further helped in elucidating the role of GQSes (Gonzalez- Pena et al., 2007; Palm et al., 2008; Wu et al., 2008).

Several studies have suggested the effect of either absence or presence of GQSes on the replication and transcription machinery, which can lead to genetic instability (Paeschke et al., 2011; Varshney et al., 2020). Several SF1B helicases, such as Pif helicases resolve these secondary structures of DNA and maintain genomic homeostasis and telomere length (Bessler et al, 2001; Li et al., 2014; Zhou et al., 2014; Byrd and Raney, 2017; Lu et al., 2018). The mutation in Pif1 helicase in *Sachharomyces* caused slow replication fork progression and breakage of chromosome near G4 motifs and aberrant chromosomal segregation (Paeschke et al., 2011; Steinacher et al., 2012). In mice, Pif1 helicase mutation accelerated weight gain and decreased locomotor activity as well as mitochondrial myopathy (Bannwarth et al., 2016; Belmonte et al., 2019). Pif mutation resulted in reduced break-induced replication (BIR) frequency at induced double-stranded breaks in mammalian cells (Li et al., 2021). These genetic studies suggested an important role of Pif helicases in various cellular functions.

Although a few studies have predicted GQSes in different plants (Garg et al, 2016; Ma et al., 2022), Pif helicases have not been characterized in plants so far. Sequence comparison allowed the identification of the ScPif1 and HsPif1 homologs in Arabidopsis (Bochman et al., 2010). However, functions of Arabidopsis Pif1-like helicase (AtPLH) have not yet been reported. In this study, we identified AtPLH genes in Arabidopsis and performed their characterization. The GQSes in one of AtPLH gene mutant in Arabidopsis (*Atplh1)* were identified at whole genome level by DNA affinity purification followed by sequencing. Several differentially bound peaks and differentially expressed genes in *Atplh1* mutants have been identified. Our study highlights the preferential genomic distribution of GQS forming regions and gene regulatory network affected in *Atplh1*.

## Results

### Identification and structural characterization of Pif-like helicases (PLH) in Arabidopsis

We identified PIF1 homologs mainly in the *Brassicaceae* and *Poaceae* families utilizing two different approaches. First, using yeast Pif1 (ScPif1) helicase protein sequence as a query in PSI-BLAST, we identified 208 homologous sequences across *Brassicaceae* and *Poaceae*. Likewise, using human Pif1 (HsPif1) helicase sequence as a query, we acquired 472 homologues. Overall, a final set of 415 non-redundant sequences was obtained (**Supplementary Table 1**).

Next, we focused on Pif1 homologs in Arabidopsis only for further characterization. In total, we identified three potential Pif1-like helicases from *Arabidopsis*, referred to as AtPLH1, AtPLH2 and AtPLH3, hereafter. These protein sequences showed higher similarity to ScPif1 (38% for AtPLH1, 46% for AtPLH2 and 47% for AtPLH3) as compared to HsPif1 (∼30%) (**Supplementary Table 2a**). A recent study suggested that Pif sequences in Arabidopsis emerged from Helitron/Helentron helicase domains (Heringer and Kuhn, 2022). The phylogenetic analysis of Pif helicase proteins from human, yeast, bacteria, Chlamydomonas, *Oryza*, *Panicum*, *Aegilops*, Arabidopsis along with TraI_A, helitron and helentron helicase domain sequences suggested that AtPLH1 is closely related to Pv_Helitron and Pif helicases. However, AtPLH2 and AtPLH3 showed divergence from yeast and human Pif sequences and more similarity to Pif sequences identified in plants (**Figure 1a**). Interestingly, AtPLH1 seems to have diverged from the above mega clade, which suggests a potential recombination event leading to the displacement and neo-functionalization of the AtPLH homologs.

**Figure 1.**
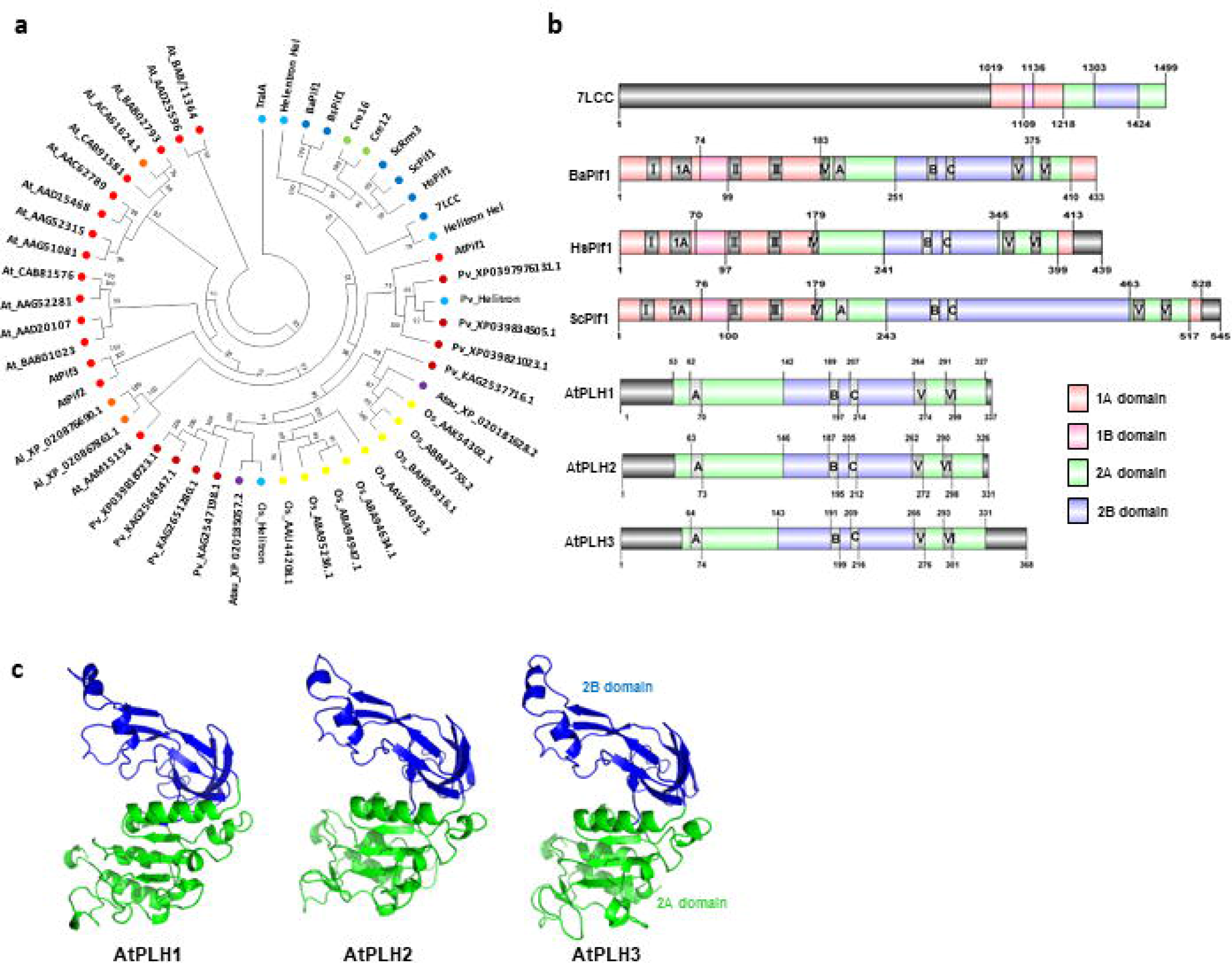
Identification of Pif Helicase homologs in Arabidopsis. a) Phylogenetic analysis of Pif helicase homologs identified from monocot and dicot plant species (Os- *Oryza sativa* subsp. *japonica*, Pv- *Panicum virgatum*, Atau- *Aegilops tauschii* subsp. *strangulata*, At- *Arabidopsis thaliana*, Al- *Arabidopsis lyrata* subsp. *petrea*) along with *Chlamydomonas reinhardtii* (Cre), Human (Hs), Bacteriodes (Ba), Bacillus (Bs) and Saccharomyces (Sc) Pif helicase sequences, Bat helitron (7LCC) and conserved Helitron and Helentron helicase domain sequences. **b)** Domain and motif organization in different Pif helicases of various organisms and Pif like helicase (AtPLH) proteins identified from Arabidopsis. **c)** Structural organization of AtPLH proteins using homology modelling, highlighting the 2A (green) and 2B (blue) domains in all three proteins.

Pif1 helicase proteins consist of three main domains: an accessory N-terminal domain, a core helicase domain and C-terminal domain. The helicase domain is further comprised of structurally conserved subdomains: 1A, 1B, 2A, 2B and 2C. All three Arabidopsis homologs, AtPLH1, AtPLH2 and AtPLH3, were found to harbor 2A and 2B subdomains (**Figure 1b**). The conserved subdomains in Pif helicases can further be identified using conserved motifs: L, L, L, L, L, L, A, B and C (Hall and Matson, 1999; Bochman et al., 2011). The motifs L and L of domain 2A and motifs A, B and C of domain 2B were identified in all three AtPLH proteins (**Figure 1b**). However, Pif1 signature motif was missing in all three AtPLH proteins, as has been reported earlier too (Bochman et al., 2010). Domain 2B forms a SH3-like domain which undergoes conformational changes upon ATP binding, allowing the protein to open or close around the DNA (Subramanya et al., 1996, Velankar et al., 1999, He et al., 2012, Chen et al., 2016). The residues present in motifs L, L, L and L are key for interacting with ATP as well as DNA (Zhou et al., 2016). Conservation of 2A and 2B domains and absence of 1A domain could potentially mean that the identified Pif1 homologs in *Arabidopsis* might have evolved from a helitron due to neofunctionalization.

The homology modelling of AtPLH proteins using helitron structure (7LCC) as a template (Kosek et al., 2021) showed structural conservation of domains 2A and 2B (**Figure 1c)**. The modelled structures were validated via Ramachandran plot (>90% in allowed region), ERRAT (>75%) and Verify 3D (>73%) (**Supplementary Table 2b)**. The structural conservation of 2A and 2B domains suggests helicase activity in AtPLH proteins to resolve GQSs.

### Genome-wide identification of putative GQS forming regions

For global identification of putative GQS-forming regions in Arabidopsis, homozygous *Atplh1* mutants (MT) from the T-DNA lines were selected (**Supplementary** Figure 1). DNA affinity purification followed by sequencing (DAP-seq) of the wild-type (WT) and *Atplh1* seedlings was performed. A total of 534 million high-quality paired-end reads for DAP-seq and input DNA samples were mapped on the Arabidopsis genome for peak calling (**Supplementary Table 3**). A total of 26053 and 30553 peaks were identified in WT and *Atplh1* mutant, respectively (**Supplementary Table 4**). The overall distribution on the Arabidopsis chromosomes showed highest peak density near the centromere regions (**Figure 2a**). A higher number of these peaks were present within the exon regions (∼41%) followed by promoter-TSS regions (21%), TTS (17.5%), intronic (10.3%) and intergenic regions (10.3%) (**Figure 2b**). The selective enrichment of GQSs in the exonic regions suggests their role in the regulation of transcriptional or translational rates. Further, these peak sequences harbored G-rich cis-regulatory elements representing the binding sites of TFs, such as BPC1 (single G repeats), ERF (double G repeats), and TRP/TRB (triple G repeats) (**Supplementary** Figure 2a).

**Figure 2.**
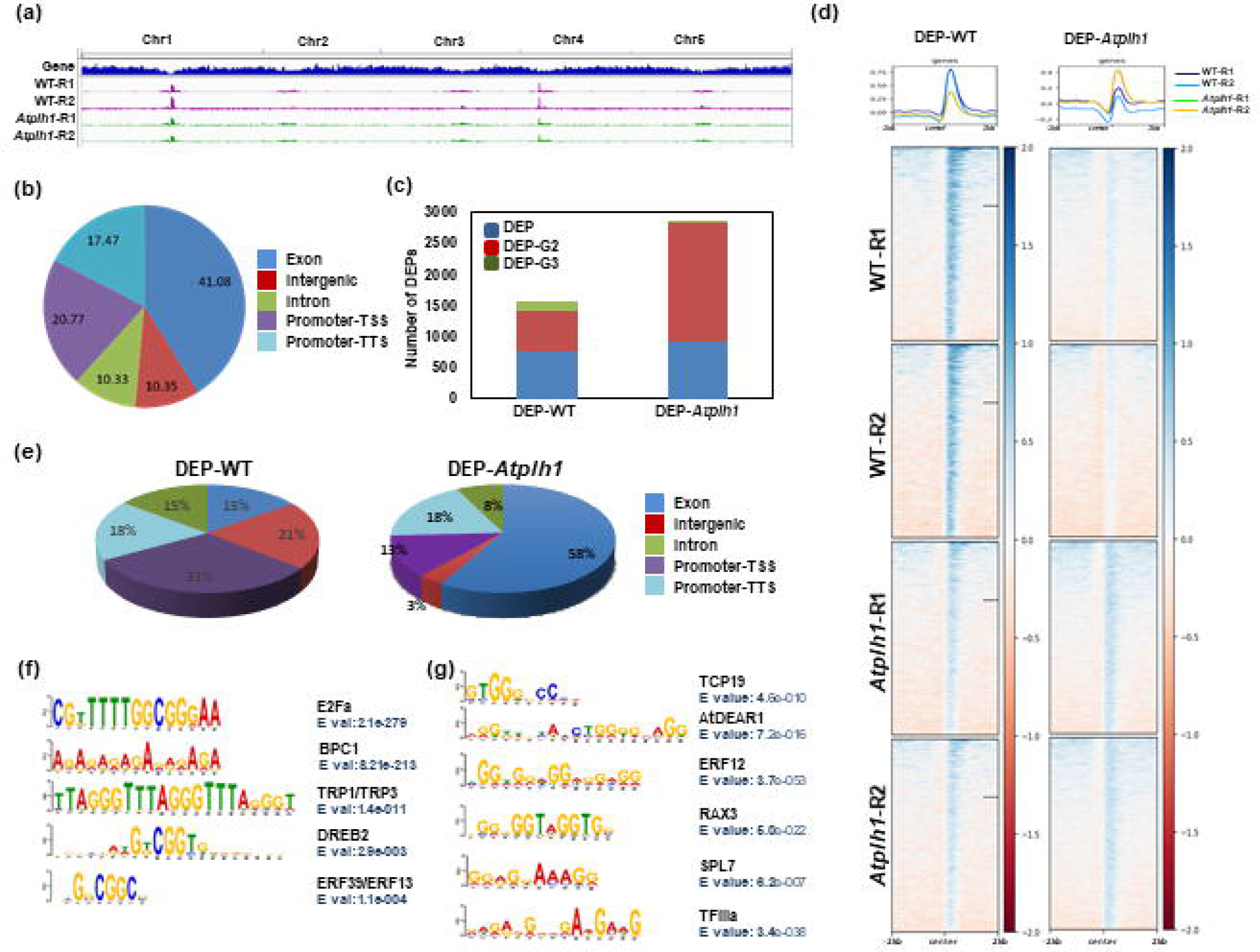
Genome wide identification of GQS binding regions using DAP-seq. a) IGV view of DAP-seq peaks in wild type (WT) and *Atplh1*. DAP- DNA affinity purified. **b)** Pie chart showing genomic region distribution of common peaks. **c)** Number of DEPs overlapping with predicted G2 and G3 sequences. DEP- Differentially Enriched peaks. **d)** Deeptools view of fragment enrichment at the differentially enriched peaks identified in WT (DEP-WT) and *Atplh1* (DEP-*Atplh1*). **e)** Pie chart representing genomic distribution of DEPs in WT and *Atplh1*. **f) & g)** Transcription factor binding motifs enriched in DEP-WT **(f)** and DEP-*Atplh1* **(g)**.

Further, we identified differentially enriched peaks (DEPs) between WT and *Atplh1* (**Figure 2c, d**). Of the 4419 DEPs identified, 1576 peaks were enriched in WT (WT-DEP) and 2843 peaks were enriched in *Atplh1* (*Atplh1*-DEP). Among these, 814 DEPs in WT and 1924 DEPs in *Atplh1* overlapped with predicted G2 or G3 type GQS forming regions in the Arabidopsis genome (**Figure 2c and Supplementary Table 4, 5**). The DEPs enriched in WT were majorly located in the promoter regions, whereas DEPs enriched in the mutant were present majorly in the exonic regions (**Figure 2e**). Several G-enriched transcription factor binding motifs, such as E2Fa, BPC1, TRP1/TRP3 and DREB2, were found enriched in the WT DEPs (**Figure 2f)**. However, binding motifs for transcription factors, TCP1, STOP1/DEAR1, TFIIIa/ERF12, RAX3, SPL7 and TFIIIa/NAC68, were enriched in the mutant DEPs (**Figure 2g**). These results suggested that the binding of transcription factors might be affected by G-quadruplex formation in these regions in the absence of AtPLH1 protein.

Chromosomal distribution of DEPs in Arabidopsis suggest high peak density in centromeric regions (**Supplementary** Figure 2b). In a recent study, 36 chromatin states based on diverse epigenetic modifications have been identified in *Arabidopsis* (Liu et al., 2018). We analysed the chromatin states of the DEPs identified in our study. The analysis revealed the enrichment of chromatin states 31, 35 and 36 in WT-DEPs, whereas chromatin states 3, 33, 34 and 36 were found enriched in *Atplh1*-DEPs (**Supplementary** Figure 3). State 36 represents the genomic regions associated with tRNA, rRNA and centromere and 1 kb promoter having CENH3, H3K9me2, and DNA methylation, while states 33 and 34 are associated with TEs and miRNAs and possess H3K9me2 and DNA methylation marks. State 3, however, is associated with coding regions and harbours H3K4me1, H3.3 and H3.1 epigenetic marks. The states 31 and 32 represent intergenic regions with DNA methylation and H3K9me2 marks, whereas state 35 is associated with intergenic and pericentromeric regions with H3K9me2, DNA methylation and H2A.x epigenetic marks. These results suggest association of the DEPs enriched in WT and *Atplh1* with different genomic regions and epigenetic marks. Further, perturbation of *AtPLH1* affects the formation of GQSes in the coding and TE regions.

### Identification of genes differentially expressed in *Atplh1* mutants

To understand AtPLH1-mediated regulation of gene expression, RNA-seq analysis of mock-treated and pyridostatin (PDS)-treated WT and *Atplh1* seedlings was performed. PDS is known to stabilize GQS structure formation *in vivo* (Rodriguez et al., 2008). The RNA-seq reads (∼688 million) obtained from different samples, WT (mock-treated WT), WT-PDS (PDS-treated WT), *Atplh1* (mock-treated *Atplh1*) and *Atplh1*-PDS (PDS-treated *Atplh1*) were mapped to the Arabidopsis genome (**Supplementary Table 6**). A total of 553 differentially expressed genes (DEGs) were identified between WT and *Atplh1* (WT vs *Atplh1*), 193 DEGs in WT after PDS treatment (WT vs WT-PDS), while 686 DEGs between WT and *Atplh1*- PDS (WT vs *Atplh1*-PDS) (**Figure 3a and Supplementary Table 7)**. Higher number of upregulated genes (425, 132 and 519) were identified as compared to downregulated genes (128, 61 and 167) in all comparisons (**Supplementary Table 7)**. A comparative analysis revealed 81 genes to be commonly regulated in WT vs *Atplh1*, WT vs WT-PDS and WT vs *Atplh1*-PDS, suggesting regulation of these genes by GQS stabilization in the presence of PDS or the absence of AtPLH1 (**Figure 3a)**. Further, 53% (294) of the DEGs are regulated in *Atplh1* seedlings in the absence or presence of PDS (WT vs *Atplh1* and WT vs *Atplh1*-PDS).

**Figure 3.**
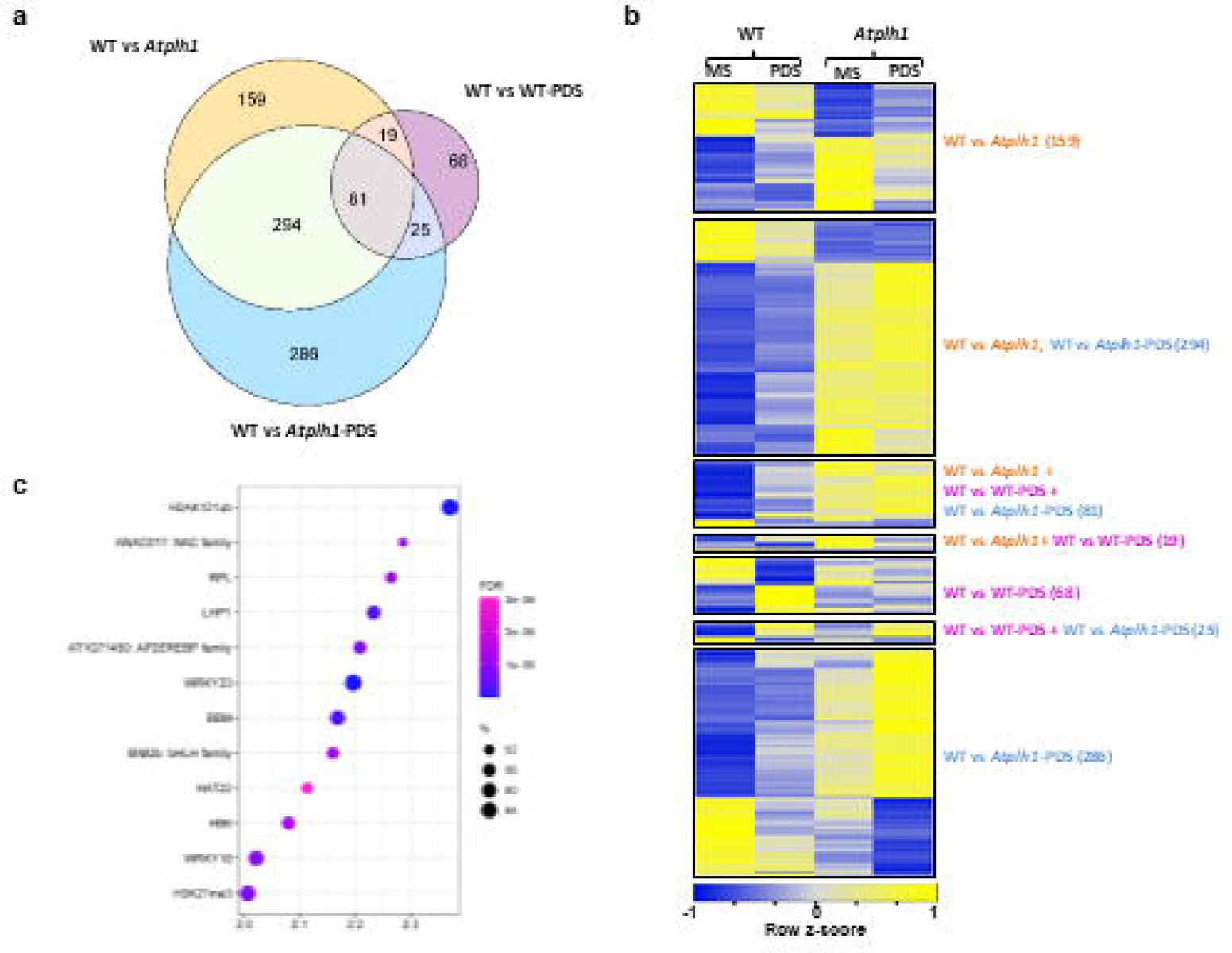
Differentially expressed genes in *Atplh1*. a) Venn plot representing number of DEGs identified in WT vs *Atplh1* (WT vs *Atplh1*), WT under control and PDS treatment (WT vs WT-PDS) and WT under control and *Atplh1* under PDS treatment (WT vs *Atplh1*-PDS). **b)** Heat map representing gene expression of DEGs. **c)** Motif enrichment of 81 genes differentially expressed in all three comparisons (WT vs *Atplh1*, WT vs *Atplh1*-PDS, WT vs WT-PDS).

Interestingly, expression of majority of the genes varied in similar direction (upregulation or downregulation) in presence of PDS and in *Atplh1* (**Figure 3b)**.

The functional annotation of 81 commonly regulated DEGS in WT vs *Atplh1*, WT vs WT-PDS and WT vs *Atplh1*-PDS suggested their involvement in processes like response to jasmonic acid signaling, response to wounding, response to osmotic and salinity and response to ABA and oxidoreductase activity (**Supplementary** Figure 4a). However, 294 DEGs regulated in WT vs *Atplh1* and WT vs *Atplh1*-PDS, were involved in iron ion starvation, floral meristem determinacy, response to wounding, salt, ABA, JA and cold regulation of transcription, protein dimerization, and UDP-glycosyl transferase activity (**Supplementary** Figure 4b). In comparison, genes regulated by PDS in WT or mutant (WT vs WT-PDS and WT vs *Atplh1*-PDS, 25 genes) were involved in pathways occurring in the extracellular region (**Supplementary** Figure 4c). RT-qPCR validation of several randomly selected DEGs suggested a very good correlation with RNA-seq based gene expression results (**Supplementary** Figure 5).

The cis-regulatory motif enrichment analysis in promoters of 81 DEGs suggested enrichment of ERF/AP2 binding sites, such as LEP, ERF55, CRF and ABR (**Supplementary** Figure 6a). Further, transcription factor enrichment analysis suggested enrichment of H3AK121ub and H3K27me3, LHP1, WRKY18, WRKY33, BBM, BIM2, RPL, ANAC017, HAT22 and HB6 (**Figure 3c)**. Recently, H2AK121ub mark has been found to be associated with less accessible, but permissive chromatin at transcriptional regulation hotspots (Yin et al., 2021; Zhang et al., 2023). The LHP1 interaction with PRC1 catalyzes H2A monoubiquitination and with PRC2 establishes H3K27me3 (Feng and Lu, 2017). The AP2/ERF transcription factor, BBM, accumulates in the quiescent center, columella stem cells and in provascular cells and mutation of *BBM* with *PLT3* causes shorter root phenotype, while its overexpression could induce somatic embryogenesis in multiple species (Galinha et al., 2007, Chen et al., 2022). The transcription factors BIM, HAT, WRKY18 and WRKY33 were shown to be involved in shade avoidance syndrome (SAS) in Arabidopsis (Cifuentes-Esquivel et al., 2013; Rosado et al., 2022). These results suggest that GQS stabilization can affect various cellular processes.

Further, motif enrichment analysis in promoters of 294 DEGs regulated in *Atplh1* mutants suggested enrichment of BPC6 and ERF binding sites (**Supplementary** Figure 6b), while transcription factor enrichment analysis suggested binding sites of bHLH18, MYB40, AGL55, MYB43, ERF55/WIND3, BAM8, WRKY18, WRKY33 and WRKY40 in these genes (**Supplementary** Figure 6c). The JA-responsive genes, bHLH18 and bHLH25, have been shown to reduce the expression of iron-uptake genes, decreasing iron acquisition from the environment (Cui et al., 2018). The transcription factor MYB40 was shown to inhibit the expression of phosphate transporter *PHT1* (Chen et al., 2021). The BAM8 protein binds to BBRE site and is involved in transcriptional activation and its overexpression resulted in smaller rosettes with rounded and hyponastic leaves (Soyk et al., 2014). ERF55/ERF58 was shown to repress genes that promote seed germination (Li et al., 2022). In addition, ERF55 or WIND3 are also involved in wound-induced plant regeneration pathway (Ikeuchi et al., 2022). The WRKY binding sites are enriched with chromatin-remodeling protein BRAHMA (BRM), near TSS and regulate various plant development processes (Guo et al., 2022). AP2/ERF and BZR transcription factors are known to regulate plant growth and development along with stress-responsive gene expression (Xie et al., 2019). MYB3 was shown to antagonize ICE1 to regulate CBF-mediated gene expression to regulate freezing stress tolerance along with role in lignin biosynthesis genes during secondary wall formation in Arabidopsis (Zheng et al., 2023, Geng et al., 2020). These results suggest that mutation in *AtPLH1* alter ERF, brassinosteriod, JA and cold regulated gene expression and plant development.

### Identification of differentially expressed genes associated with GQSes

To study the effect of GQSs in mediating differential gene expression, we identified the DEPs associated with DEGs. A total of 21 DEGs overlapped with WT-DEPs and 41 DEGs overlapped with *Atplh1*-DEPs, of which three DEGs harbor DEPs enriched in both WT and *Atplh1* (**Figure 4a and Supplementary Table 8**). The majority of these DEGs harbor DEPs in the exonic regions (27 and 8 DEGs in *Atplh1-*DEP and WT-DEP, respectively), while few DEGs harbor DEPs in the promoter region (3 and 6 DEGs in *Atplh1-*DEP and WT-DEP, respectively). Gene ontology enrichment analysis of the DEP associated DEGs (WT-DEP- DEG) revealed enrichment of pathways like protein self-association, response to wounding, and response to salt stress (**Figure 4b**). However, DEGs associated with DEP enriched in mutants (*Atplh1-*DEP-DEG) were involved in functions like unfolded protein response, quercitine-7-O-glucosyltransferase activity, quercitine-3-O-glucosyltransferase activity, and response to heat response (**Figure 4c**).

**Figure 4.**
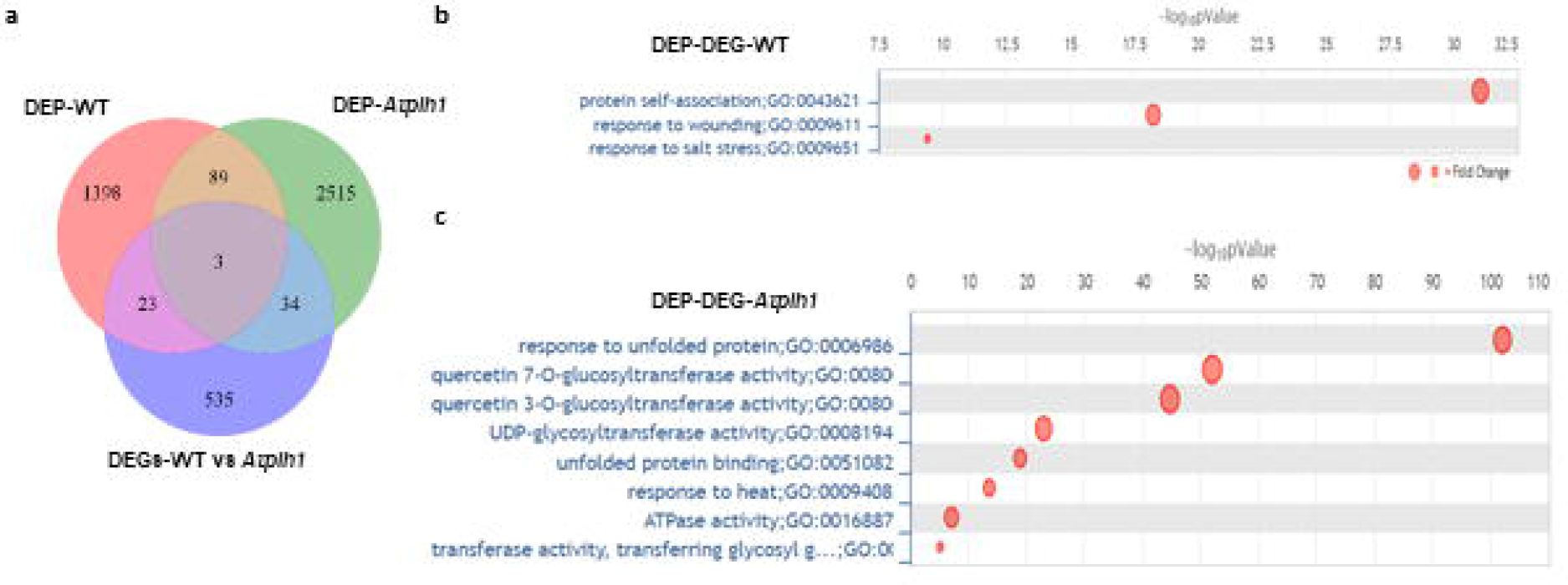
Identification of GQS regions regulating gene expression. a) Venn plot representing overlap of DEGs (WT vs *Atplh1*) with DEP-WT and DEP-*Atplh1*. **b)** and **c)** Bubble plot representing GO enrichment significance of the GO terms in DEP-DEGs identified in WT **(b)** and *Atplh1* **(c)**.

Further, GQS formation in the randomly selected DEP-DEGs was validated by *in-vitro* assays including CD-spectroscopy and ChIP-qPCR. These included DEGs (UDP-glucosyl transferase 85A3, UGT85A3; AT-Hook Motif Nuclear Localized Protein 17, AHL17; Phospholipase D Beta 2, PLD Beta 2; Dark Inducible 11, DIN11) that harbored DEPs within gene body regions (**Supplementary** Figure 7). The CD spectroscopy analysis suggested parallel GQS formation (negative peak at 240 nm and positive peak around 260 nm) in the presence of K^+^ in all four sequences (**Figure 5a**). Further, ChIP-qPCR for all four DEPs, confirmed enrichment of GQSs in WT for DIN11, while remaining three regions showed enrichment of GQS formation in *Atplh1* (**Figure 5b**). *UGT85A3* gene is involved in root development and secondary metabolic process. AHL17 is involved in cell wall biogenesis and defense response, PLD Beta 2 has calcium ion binding and phospholipase activity and *DIN11* expression is induced upon dark treatment. These results suggest that GQS structure stabilization within these genes in *Atplh1* mutant potentially regulates gene expression and root development.

**Figure 5.**
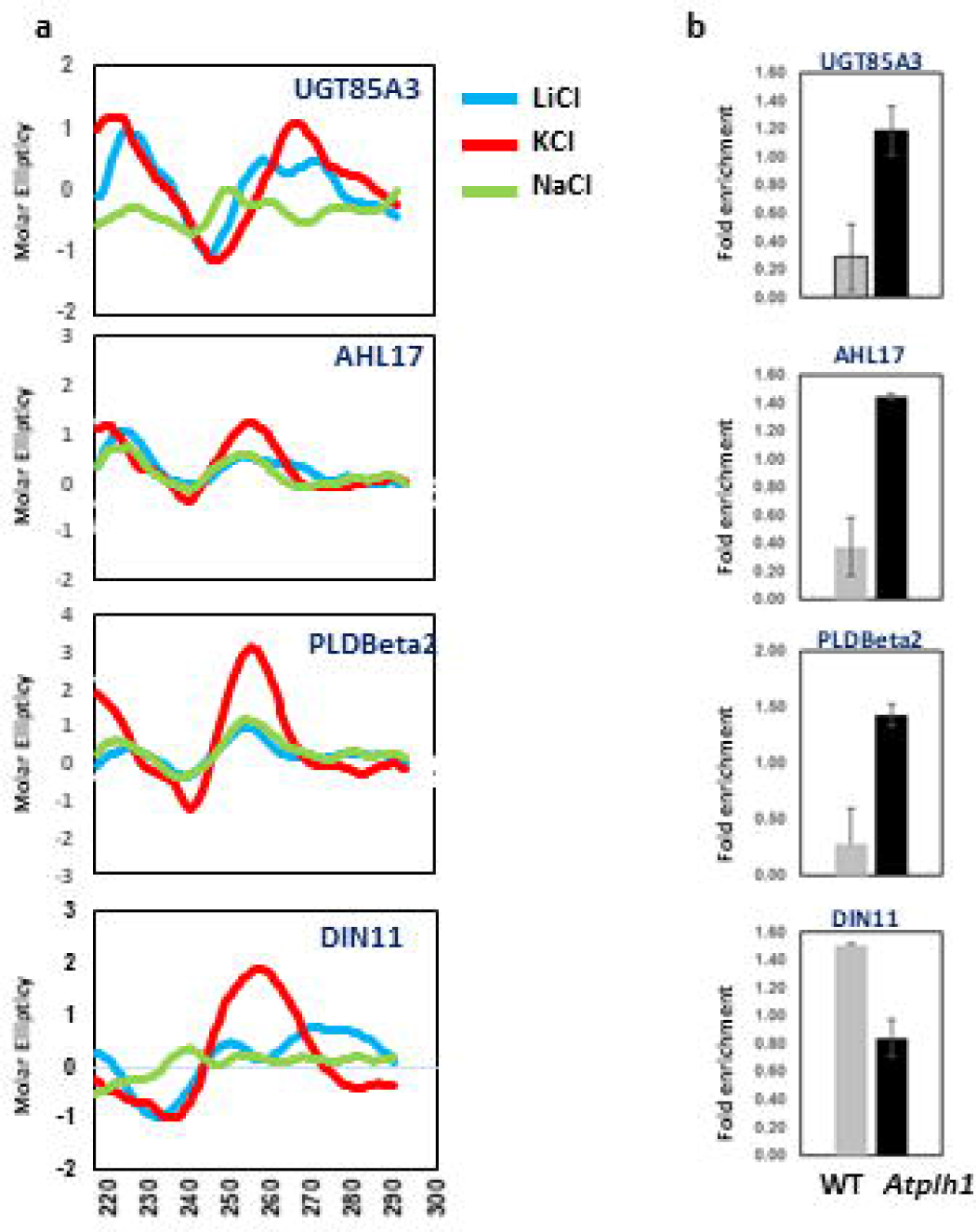
Validation of GQS formation in selected regions identified by DAP-seq. a) CD spectroscopy of selected peaks identified by DAP-seq for formation of GQS structure in different ionic conditions. **b)** Fold enrichment of selected genomic regions by ChIP-qPCR in WT and *Atplh1* mutant in DAP-purified samples.

### Cold-stress tolerance of *Atplh1* mutants

Differential gene expression analysis suggested that MYB43 target genes are upregulated in *Atplh1*. Further, MYB43 TF regulate freezing stress tolerance by antagonizing ICE1 transcription factor to regulate CBF gene expression in Arabidopsis (Zheng et al., 2023). Thus, we analyzed the response of *Atplh1* mutants under cold stress at different temperatures (4°C, 1°C and -20°C). The results suggested a lower growth inhibition in *Atplh1* as compared to WT (**Figure 6a and 6b)**. The difference is more pronounced at mild cold stress (4°C) as compared to severe cold (1°C) and freezing stress (-20°C). These results suggest that stabilization of GQSs in *Atplh1* mutants under cold temperature could result in the upregulation of downstream cold-responsive genes, and thereby promoting cold tolerance in *Atplh1*. The observed effect could be due to either the inability of MYB43 TF to bind its target genes or increased binding of ICE1 TF at the promoters of cold stress-responsive genes and hence regulation of stress response.

**Figure 6.**
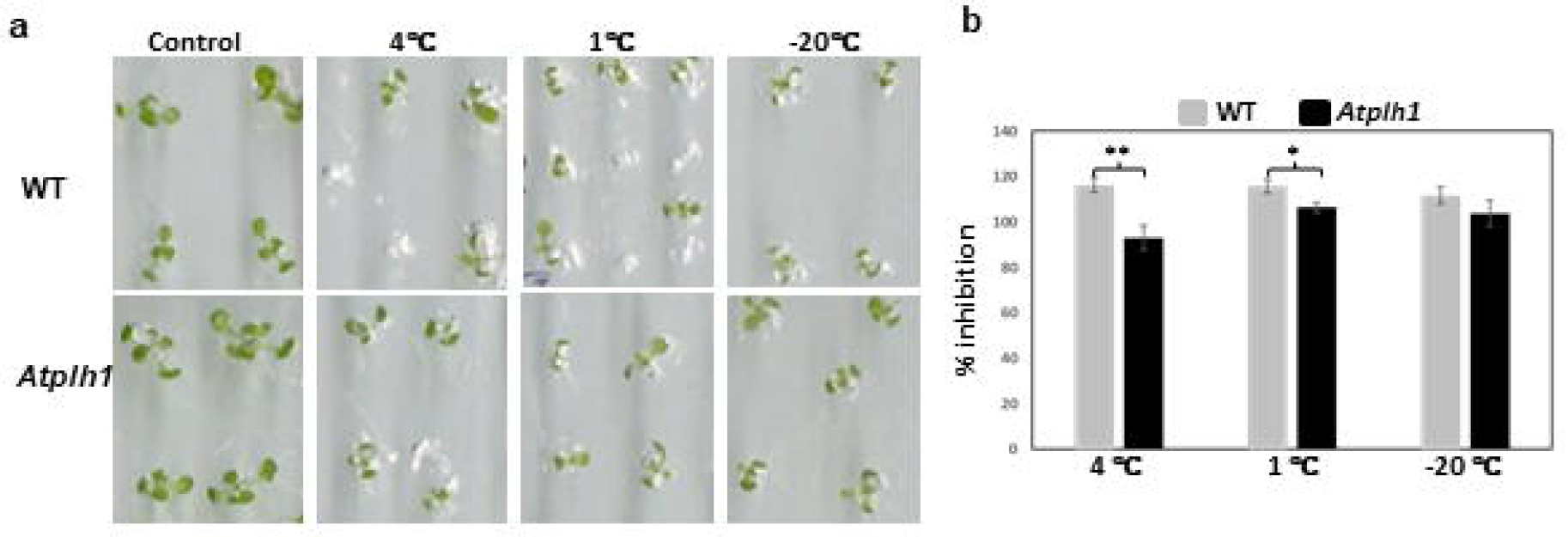
*Atplh1* mutants are tolerant to cold stress. a) Five-day old seedlings WT and *Atplh1* seedlings subjected to cold stress at 1 °C & 4 °C for thirty minutes and -20 °C for one minute. Seedlings were transferred later to 4°C for ten days and were scanned on tenth day of stress. **b)** Bar plot shows percentage inhibition in WT and *Atplh1* mutants under cold stress (**, p-value ≤ 0.05; *, p-value ≤ 0.01).

## Conclusions

We identified homologs of human and yeast Pif1 helicase in different plant species. The sequence and structure analysis of three Arabidopsis homologs revealed the presence of conserved sequence and structure of 2A and 2B subdomains suggesting a functional helicase catalytic center. The GQS-forming regions were identified in *Atplh1* mutants at global level and revealed their overlap with G1, G2 and G3 rich, transcription factor binding motif sequences. The DEPs between WT and *Atplh1* were associated with three chromatin states, including tRNA, rRNA and centromeric regions (associated with CENH3, H3K9me2 and DNA methylation), TEs and miRNAs (associated with H3K9me2 and DNA methylation) and coding regions (associated with H3K4me1, H3.3 and H3.1 epigenetic marks). This observation suggested, perturbation of *AtPLH1* could be affecting GQS formation in these regions. Subsequently, a set of 81 DEGs regulated in *Atplh1,* as well as by PDS treatment were identified. These genes were involved in processes like response to wounding, jasmonic acid signaling, response to osmotic and salinity and response to ABA and oxidoreductase activity, with TF sites for PRC1 mediated histone ubiquitinylation, WRKY and AP2/ERF TFs. However, DEGs regulated in *Atplh1* mutant (294) were targets of bHLH18, MYB43, WRKY and AP2/EREBP TFs which are involved in shade avoidance, cold-stress and wound induced regeneration response suggesting regulation of these processes in *Atplh1*. These DEGs in turn could regulate plant response to light, wounding and abiotic stress. Overall, our study identified GQS enriched regions and altered gene expression in *Atplh1.* Altered gene expression could be a result of altered binding of transcription factors or regulation of transcription through R-loop formation (which leads to G4 folding that stabilizes R-loop, Lee et al., 2020). In future, it would be interesting to study the effect of GQS stabilization on transcription factor binding or R-loop mediated transcriptional control.

## Materials and methods

### Identification of Pif helicase homologs, motif identification and phylogenetic analysis and protein modelling

Pif1 sequences were identified using PSI-BLAST with ScPif1 and HsPif1 at default parameters. Sequences matching to the Pif1 profile were searched on the PFAM database. Further, PSI-BLAST and PFAM database search was restricted to the *Brassicaceae* and *Poaceae* families. Motif analysis was carried out using Multiple Sequence Alignment (MSA) of the candidate sequences using the MEGA-X tool. The motifs were manually mapped and identified using consensus sequences from literature. MUSCLE algorithm was used to generate the alignment. The phylogenetic tree was inferred using the Maximum Likelihood method with 1000 bootstraps utilizing WAG (Whelan and Goldman) model with Frequency (F, frequency of amino acids in the dataset) and Hetrogeneity (G, gamma distribution) being taken into account. Tree was generated using IQ tree tool. Structural homology modeling of the proteins was done using Swiss-model tool based on homology with already resolved structure 7LCC, available on PDB database. The selected model structures obtained from these different approaches were validated using ERRAT, Verify3D and PROCHECK tools.

### Plant growth conditions

Wild type plants (Col-0) and Pif helicase mutant (*Atplh1*) in Col-0 ecotype background were used for this study. T-DNA insertion mutants (Salk114318.50.30x for *Atplh1*) were obtained from the Arabidopsis Biological Resource Center (ABRC). For selecting homozygous mutants from T-DNA lines for *AtPLH1;* T-DNA genotyping PCR was performed using genomic DNA of T3 lines with LB1.3, LP and RP primers (**Supplementary** Figure 1 **and Supplementary Table 9)**. Gene expression was checked by RT-PCR from seedling and flower tissue of wild type and mutant plants using gene specific primer. The seeds were surface sterilized (with 2% sodium hypochlorite, 0.1% Tween 20) and plated onto solid MS medium supplemented with 1% sucrose (w/v) followed by vernalization for 4 days under dark conditions at 4°C. The plates were then shifted to growth chamber maintained at 22°C, 14h light and 10h dark. For cold stress treatment, 5-day old seedlings were then transferred under either normal conditions or at 1°C, 4°C for 30 minutes and -20°C for one minute followed by growth at 4°C for another 10 days. Rosette area measurements were performed on 15-day old seedlings. All the experiments were done in triplicates. The measurements were done using ImageJ software.

### DNA Affinity Purification- sequencing and data analysis

Genomic DNA was isolated from ten days old Wild type (WT) and *Atplh1* mutant seedlings. Sonicated DNA was used for immunoprecipitation with BG4 antibody (anti-GQS, Merck Millipore) bound on anti-FLAG dyna beads (Invitrogen). Eluted DNA was purified using a Minelute PCR purification kit (Qiagen) and sent for sequencing. A 100 bp Illumina paired end library was generated. NGS QC Tool kit was used for quality control, followed by mapping of high- quality reads onto the *Arabidopsis* genome (TAIR10) using the Bowtie2 tool. Peak calling was done using the MACS2 tool (p-value- 0.05) and peaks were identified with respect to input control (sheared DNA) and annotated using the HOMER tool. BEDtools was used to overlap the peaks identified to the previously reported G_2_L_1-4_ and G_3_L_1-7_ GQS motifs in Arabidopsis (Garg et al., 2016). Differentially enriched peaks (DEPs) were identified using the Diffbind tool (cutoff of 0.8 FC and p value 0.05) between WT and *Atplh1.* Motif enrichment analysis for peak regions was done using 250 bp upstream and downstream sequence from peak center using MEME-ChIP.

### RNA sequencing and analysis

WT and At*plh1* mutant seedlings were grown on MS-medium plates under standard conditions for 5 days, then transfered to MS-medium with/without GQS stabilizing ligand Pyridostatin (PDS) and grown for another 5 days. The 10-day old seedlings were then used to isolate total RNA and samples were sent for paired end sequencing (100 bp read length). High-quality reads were mapped onto the Arabidopsis genome (TAIR10) using HISAT2. The transcripts were constructed through Stringtie and the differential expression was then determined using Cuffdiff (fold change ≤ 2 and *P*-value ≤ 0.05). Gene ontology (GO) enrichment analysis, motif analysis and transcription factor binding enrichment analysis of the differentially expressed genes was performed using Plant Regulomics (http://bioinfo.sibs.ac.cn/plant-regulomics/). BEDtools intersect was used to identify DBPs present in either gene body or 2kb upstream of the identified differentially expressed genes (DEGs). Heatmaps using the FPKM values were generated using Heatmapper online tool.

### Quantitative Real Time- Polymerase Chain Reaction

The cDNA samples were synthesized from 2 µg of RNA using a High-capacity cDNA synthesis kit (Applied Biosystems). RT-qPCR was performed using the ABI SYBR green kit as per manufacturer’s recommendation. For qPCR validation of DEPs, primers were designed from the flanking sequence from the center of the peak (±125 bp). ChIP-qPCR was performed using both Input and ChIP-DNA samples along with PP2A as a non-target region. For each primer set three technical replicates and two biological replicates were set up.

### Circular Dichroism Spectroscopy

Oligos of 30-35 bp length were obtained from IDT (**Supplementary Table 9**). Oligo structures were prepared in KCl, NaCl and LiCl ionic conditions as mentioned earlier (Singh et al., 2020). 10 µM of oligos along with 150 mM of NaCl/KCl/LiCl were analyzed using a CD-spectrophotometer (Applied PhotoPhysics Chirascan).

## Author Contributions

RG: Conceptualization; SS, SS, HS performed wet lab experiments, KG and MS: Data curation and NGS data analysis; SB: Homologous protein identification and analysis; RG: Funding acquisition, Project administration; Resources, Supervision; RG, SS, SS, and KG: writing, review & editing.

## Acknowledgments

SS, SS, HS and KG acknowledges research fellowship from Shiv Nadar Institution of Eminence (Deemed to be University). The authors acknowledge funding support by SERB ECRA and Shiv Nadar Institution of Eminence (Deemed to be University). We are thankful to Dr Hardik Gala and Prof Mukesh Jain for insightful comments and suggestions on the manuscript.

## Conflicts of Interests

The author declares no competing interests.

## Data availability

DAP sequencing data and RNA sequencing data have been submitted to NCBI’s Gene Expression Omnibus (GEO) with the accession number GSE250104.

## Funding

The work was financially supported by SERB ECRA and Shiv Nadar Institution of Eminence.

## Notes

### Competing Interest Statement

The authors have declared no competing interest.

