## Supplementary material for "Genome-Wide Identification of G-Quadruplex forming regions in Arabidopsis: Unraveling the Role of Pif like Helicase (AtPLH1) in Gene Regulation and Stress Response": Supplementary-Figures.pdf

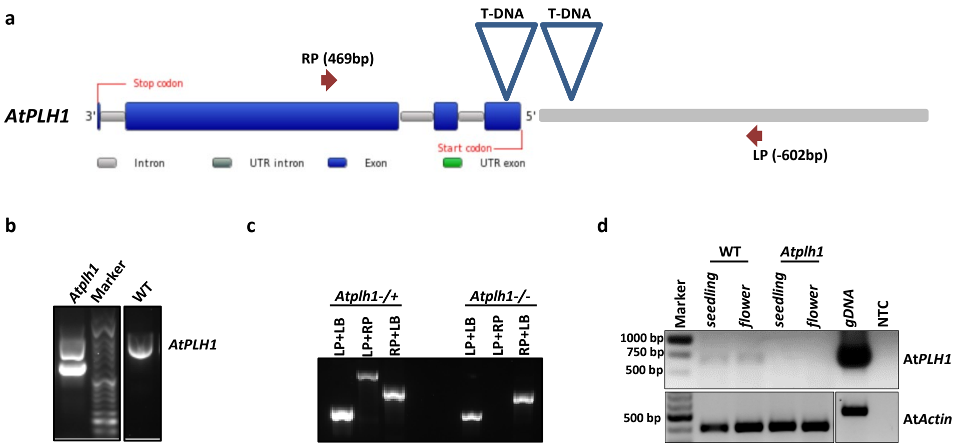


**Supplementary Figure 1**. **(a)** Schematic representation of T-DNA insertion lines for *AtPLH1* with position of left primer (LP) and right primer (RP) specified in brackets along with primers. **(b)** Genotyping PCR for homozygous lines of *Atplh1* mutant along with Col0 (WT) using LP and RP primers specific for *AtPLH1* along with LB1.3 primer (LB). **(c)** Genotyping PCR for heterozygous and homozygous lines of At*plh1* mutant using LP, RP primers specific for *AtPLH1* along with LB1.3 primer. **(d)** RT-PCR for *AtPLH1 and AtActin* gene from seedling and flower RNA from wild type (WT) and *Atplh1* mutant. gDNA: genomic DNA, NTC: No template control.

**a**

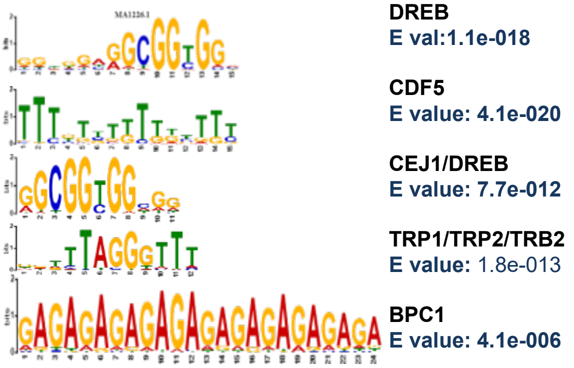


**b**

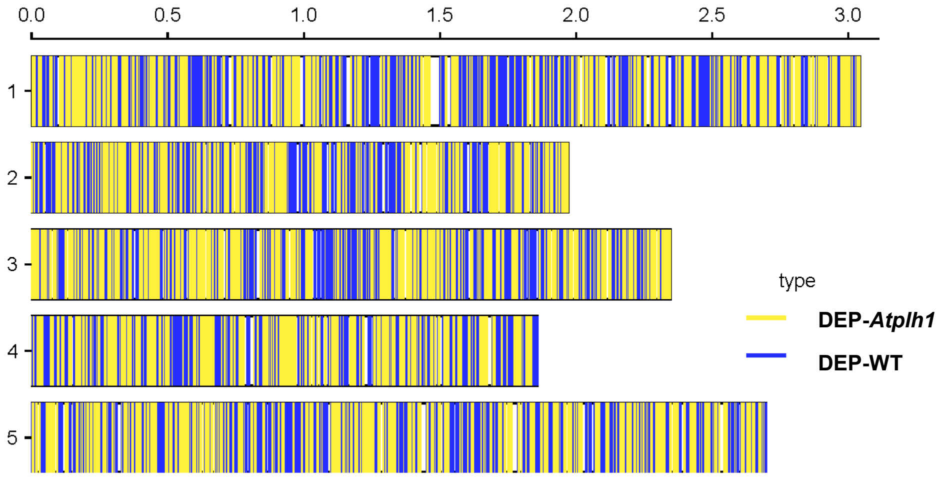


**Supplementary Figure 2**. **(a)** Motif enrichment by MEME in 1 kb sequence centered on commonly identified peak regions (only motifs centrally enriched are shown). **(b)** Chromosomal distribution of identified DEPs.

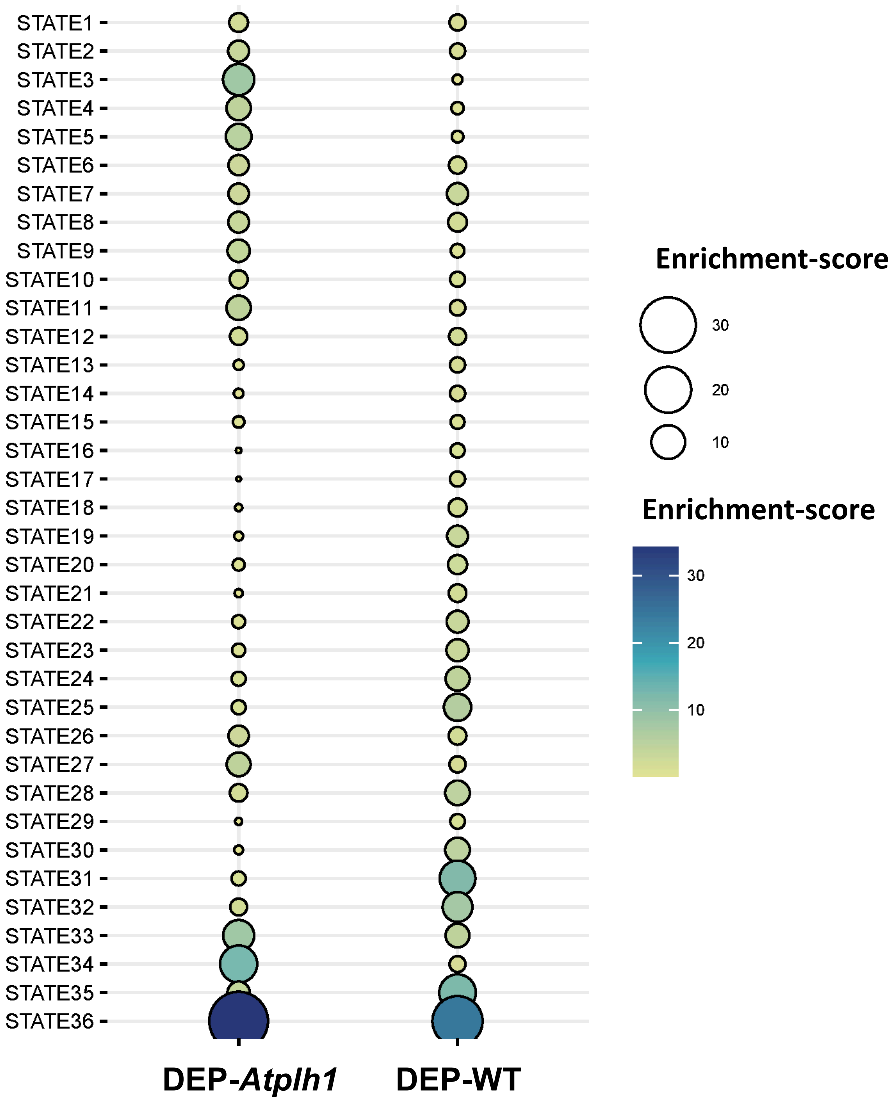


**Supplementary Figure 3.** Enrichment score of peaks enriched in WT (DEP-WT) and *Atplh1* (DEP-*Atplh1*) in 36 different chromatin states identified in Arabidopsis associated with different chromatin features and regions in the genome identified in Liu et al., 2018.

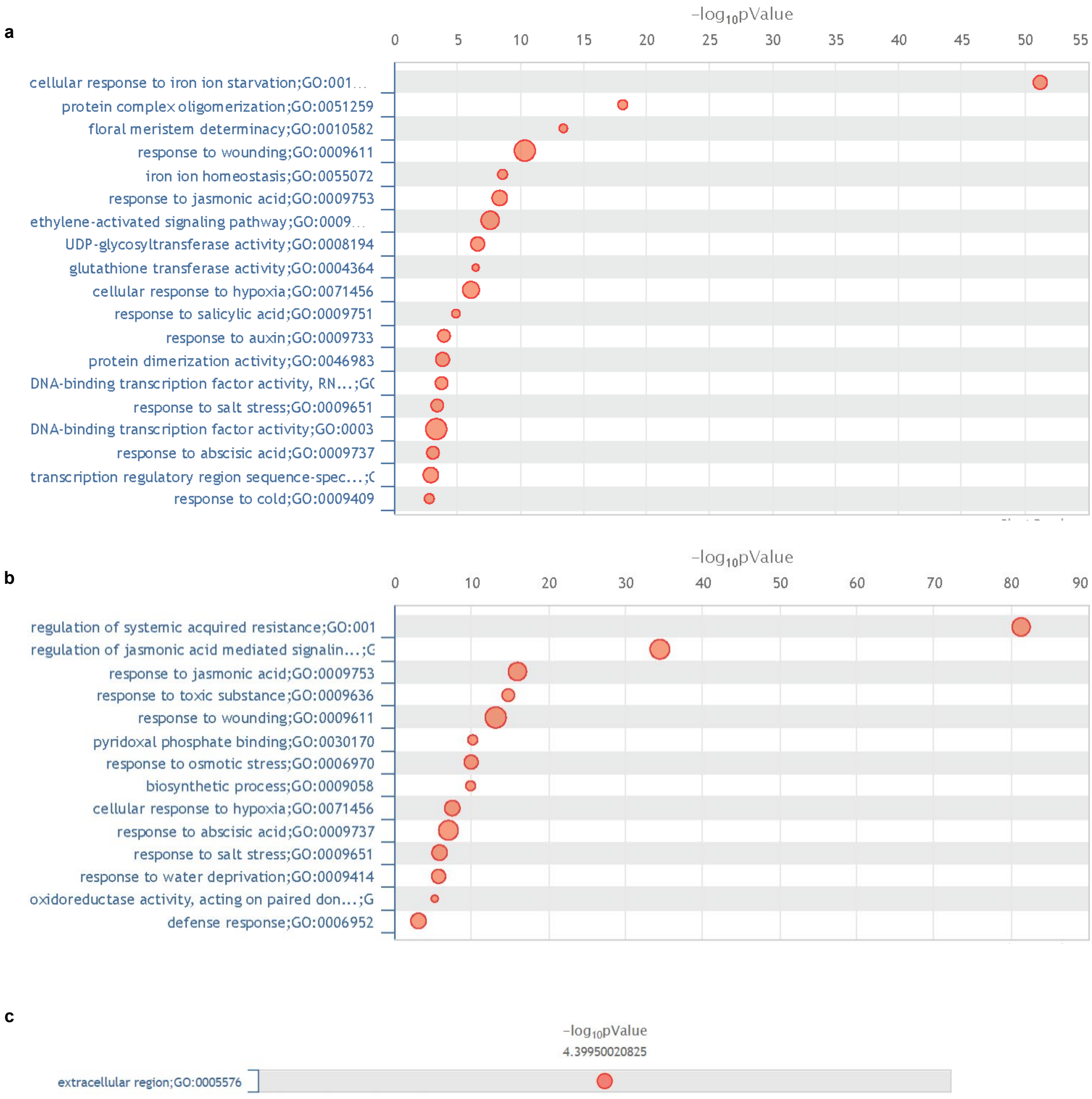


**Supplementary Figure 4**. GO enrichment analysis of **(a)** 294 DEGs common between WT vs *Atplh1* and WT vs *Atplh1*-PDS, **(b)** 81 DEGs common between WT vs *Atplh1,* WT vs *Atplh1*-PDS and WT vs WT-PDS comparison, **(c)** 25 DEGs common between WT vs WT-PDS and WT vs *Atplh1*-PDS. Size of each circle represents fold change value and x-axis represents enrichment significance p-value.

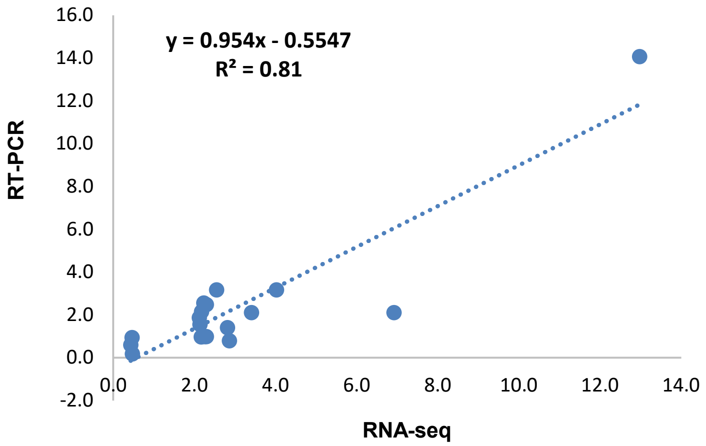


**Supplementary Figure 5.** Correlation of qRT-PCR and RNA-seq fold change for a set of selected genes.

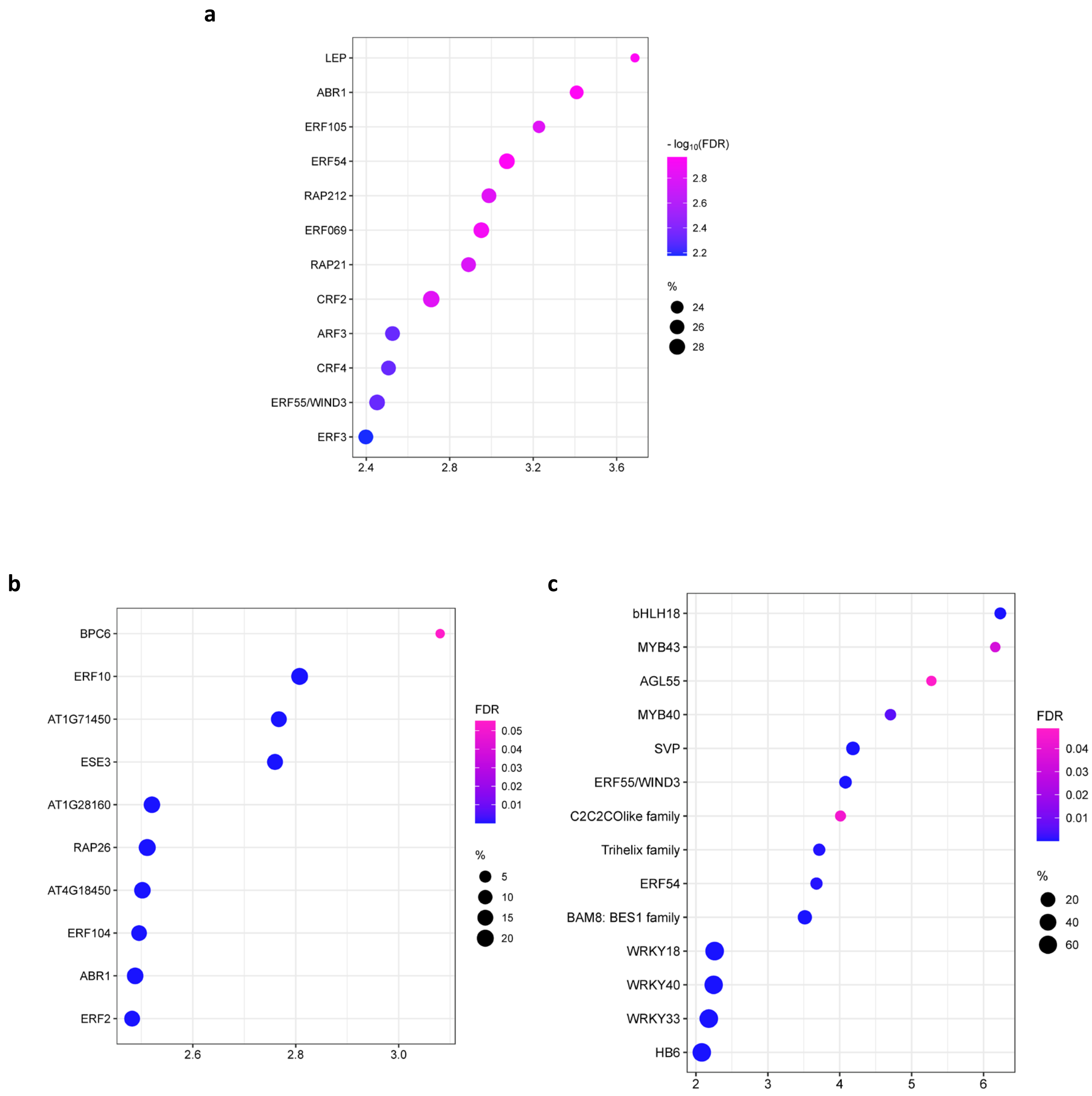


**Supplementary Figure 6**. **(a)** Motif enrichment of 81 DEGs common between WT vs *Atplh1* and WT vs *Atplh1*-PDS. **(b)** Motif enrichment analysis of 294 DEGs (WT vs *Atplh1*, WT vs *Atplh1*-PDS and WT vs WT-PDS) and **(c)** Transcription factor enrichment analysis of 294 DEGs (WT vs *Atplh1*, WT vs *Atplh1*-PDS and WT vs WT-PDS).

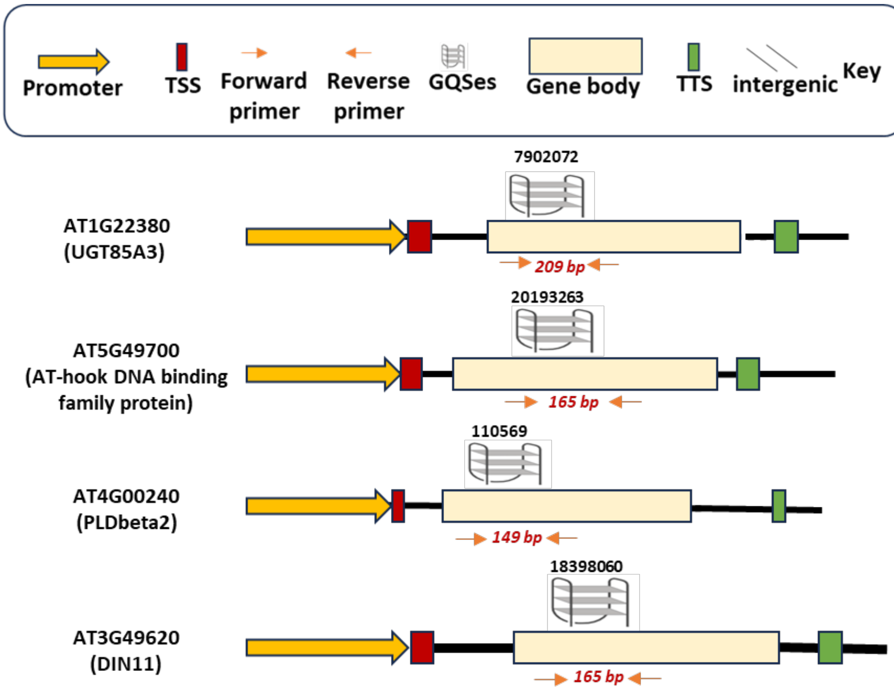


**Supplementary Figure 7**. Diagrammatic representation of DBPs validated through Chip-qPCR depicting their central position above the GQS structure along with its location within the gene and size of PCR products depicted by red font flanked by orange arrows within the peak regions.
